# Radionuclide selection influences imaging outcomes in immunoPET with a brain-penetrant anti-Aβ antibody

**DOI:** 10.1101/2025.05.31.657140

**Authors:** Sara Lopes van den Broek, Klas Bratteby, Ximena Aguilar, Thuy A. Tran, Stina Syvänen, Dag Sehlin

## Abstract

**Background:** Bispecific antibodies exploiting receptor-mediated transcytosis offer a promising strategy to overcome limited blood-brain barrier permeability in Alzheimer’s disease (AD) therapy and imaging. Lecanemab-Fab8D3 (Lec-Fab8D3), a bispecific anti-amyloid beta (Aβ) antibody engineered for enhanced brain delivery, holds potential as a companion immunoPET imaging diagnostic with the novel lecanemab immunotherapy. This study aimed to compare three radionuclides—zirconium-89 (^89^Zr), copper-64 (^64^Cu), and iodine-124 (^124^I)—for PET imaging with Lec-Fab8D3 to study its in vivo brain distribution and evaluate its potential as an AD companion diagnostic.

**Methods:** Lec-Fab8D3 was conjugated to DFO* or NODAGA for ^89^Zr and ^64^Cu radiolabeling, respectively, or directly radioiodinated with ^124^I. PET imaging was performed in the Tg-ArcSwe mouse model of Aβ pathology and wild-type (WT) littermates at multiple time points post administration of the radiolabeled antibody, followed by ex vivo biodistribution, autoradiography, and Aβ quantification to assess brain uptake, specificity, and distribution of the radiolabeled Lec-Fab8D3.

**Results:** Radiolabeled Lec-Fab8D3 variants showed retained binding properties with high radiochemical purity and yields. PET imaging demonstrated cortical brain uptake of all three tradiotracers in Tg-ArcSwe mice, with [^89^Zr]Zr-DFO*-Lec-Fab8D3 and [^124^I]I-Lec-Fab8D3 showing the best discrimination between Tg-ArcSwe and WT mice at 48–72 h post-injection. The highest absolute brain retention, combined with a lower brain-to-cerebellum ratio, was observed in both Tg-ArcSwe and WT mice that received the radiometal-labeled (^89^Zr and ^64^Cu) antibody, likely due to the residualizing nature of radiometals. Ex vivo analyses confirmed PET findings, and immunostaining demonstrated co-localization of Lec-Fab8D3 with Aβ deposits.

**Conclusions:** ImmunoPET imaging with bispecific Lec-Fab8D3 enables specific detection of brain Aβ pathology in an AD mouse model. ^89^Zr was superior to ^64^Cu due to a more compatible half-life, while ^124^I displayed higher regional contrast than both radiometals, despite lower overall brain signal. The combined findings from radiometal- and iodine-based immunoPET will enhance our understanding of intra-brain distribution of bispecific antibodies. Furthermore, this highlights the importance of the choice of radiolabeling strategy and how it will impact the outcome of immunoPET with bispecific Aβ antibodies.

## Introduction

Positron emission tomography (PET) is a powerful molecular imaging modality that enables non-invasive, in vivo visualization and quantification of pathological processes. In Alzheimer’s disease (AD), PET imaging has played a critical role in detecting amyloid pathology using radiotracers such as carbon-11-labeled Pittsburgh Compound-B ([^11^C]PiB) and fluorine-18-labeled radiotracers like florbetaben, flutemetamol, and florbetapir.^1–6^ These radiotracers have been instrumental in clinical trials by supporting both patient selection and therapeutic monitoring for antibodies such as lecanemab, the first approved disease-modifying treatment for AD.^7^ However, these radiotracers do not bind amyloid-beta (Aβ) directly but rather to β-sheet-rich fibrillar structures in dense-core plaques. This results in limited specificity and a potential mismatch with therapeutic targets, particularly soluble and diffuse Aβ aggregates recognized by monoclonal antibodies such as lecanemab.^8–10^

ImmunoPET, which involves radiolabeled antibodies as PET tracers, addresses this challenge by combining the high target specificity of antibodies with the sensitivity of PET imaging. However, as regular antibodies are large molecules with limited capacity to cross the blood-brain barrier (BBB), they must be redesigned to achieve efficient brain delivery. One approach is to engineer antibodies into bispecific formats that enable receptor-mediated transcytosis across the BBB. This can be accomplished by designing antibodies with dual specificity, such as combining Aβ targeting with affinity for the transferrin receptor (TfR).^11–14^ The TfR facilitates transport from the luminal side of the BBB into the brain parenchyma. Additionally, engineering strategies such as the introduction of LALA-PG mutations to the antibody’s Fc domain can minimize Fc gamma receptor (FcγR) binding, thereby reducing off-target effects such as antibody-dependent cellular cytotoxicity (ADCC).^15^ This is particularly important for diagnostic radiotracers where no pharmacological effect is desired.^16,17^

In addition to detecting Aβ pathology, PET imaging with radiolabeled bispecific antibodies offers critical insights into antibody distribution within the brain, target engagement, and therapeutic efficacy.^18,19^ This may be especially important given the development of next-generation therapeutic antibodies such as trontinemab, which utilize a bispecific design for TfR-mediated BBB transcytosis.20,21 Thus, PET with bispecific antibody-based radiotracers not only allow detection of Aβ species beyond dense-core plaques but also structurally resemble emerging bispecific therapeutic antibodies. As such, they have the potential to be developed as companion diagnostics that directly support therapeutic strategies.

To ensure successful immunoPET applications and to understand where antibodies localize in the brain, selecting an appropriate radionuclide is essential. The physical half-life of the radionuclide must align with the biological half-life of the antibody, while also offering optimal imaging properties. The most commonly used PET radionuclides with appropriate half-lives for antibody include zirconium-89 (^89^Zr), T_1/2_ ∼ 78 hrs, copper-64 (^64^Cu) T_1/2_ ∼ 13 hrs, and iodine-124 (^124^I) T_1/2_ ∼ 100 hrs, each offering distinct characteristics that influence their suitability for specific imaging scenarios (Table 1).^14,22–24^ In this study, we investigated how a bispecific variant of lecanemab, Lec-Fab8D3, radiolabeled with ^89^Zr, ^64^Cu, or ^124^I, performs as an Aβ-targeted immunoPET radiotracer in a mouse model of Aβ pathology. Lec-Fab8D3 was engineered for TfR-mediated BBB transport through incorporation of the Fab8D3 antibody fragment. We assessed how the choice of radionuclide influences the PET signal, which in turn affects the apparent distribution pattern of Lec-Fab8D3. Our findings also provide important insights into antibody biodistribution in the brain and lay the foundation for establishing radiolabeled Lec-Fab8D3 as a companion diagnostic in AD.

**Table 1:** Nuclear properties, advantages and limitations of ^89^Zr, ^64^Cu and ^124^I.

| Radionuclide | Half-Life | Decay mode | Positron Energy | Advantages | Limitations |
| --- | --- | --- | --- | --- | --- |
| Zirconium-89 ($^{89}\text{Zr}$) | 78.4 hours (3.3 days) | $\beta^+$ (23%), EC | 0.897 MeV | Long half-life matches antibody kinetics, relatively stable chelation with desferrioxamine (DFO*) | High-energy $\gamma$ emission increases radiation dose, requires chelation |
| Copper-64 ($^{64}\text{Cu}$) | 12.7 hours | $\beta^+$ (17.9%), $\beta^-$ (39%), EC | 0.653 MeV | Short half-life allows earlier imaging, theranostic potential | Moderate half-life may not fully match antibody kinetics, requires chelation |
| Iodine-124 ($^{124}\text{I}$) | 100.2 hours (4.2 days) | $\beta^+$ (23%), EC | 2.138 MeV | Long half-life matches antibody kinetics, direct labeling without chelator | High positron energy reduces resolution, limited availability |

In this study, we investigate how a bispecific variant of lecanemab, Lec-Fab8D3, radiolabeled with ^89^Zr, ^64^Cu, or ^124^I, performs as an Aβ immunoPET radiotracer in a mouse model of Aβ pathology. Lec-Fab8D3 was engineered for transferrin receptor (TfR)-mediated transport across BBB by incorporating the TfR-targeting antibody fragment Fab8D3. Additionally, Lec-Fab8D3 included LALA-PG mutations to abolish Fc gamma receptor (FcγR) binding, thereby preventing antibody-dependent cellular cytotoxicity (ADCC) and minimizing off-target interactions.^16,17,25^ This study aims to determine how the choice of radionuclide influences immunoPET detection of the bispecific antibody in the brain. The findings will provide insights into the brain distribution of Lec-Fab8D3, with important implications for therapeutic development. Moreover, this work lays the foundation for establishing Lec-Fab8D3 as a companion diagnostic in Alzheimer’s disease, enabling more precise patient stratification, real-time monitoring of treatment efficacy, and more efficient design of future clinical trials.

## Results

### Lec-Fab8D3 maintains binding affinity after conjugation and radiolabeling

Lec-Fab8D3 was successfully conjugated with a 7-fold molar excess of DFO*-NCS for radiolabeling with [^89^Zr]Zr-oxalate or with a 10-fold molar excess of p-NCS-benzyl-NODAGA-benzyl-p-NCS for radiolabeling with [^64^Cu]CuCl2. The integrity and binding capacity of the antibody after conjugation and radiolabeling were assessed by ELISA, confirming no or minimal alteration in binding to both Aβ protofibrils and the murine TfR (mTfR) (Figure 1). Molar ratios were determined based on titration experiments with varying chelator-to-antibody ratios where the highest degree of conjugation without significant loss in binding was selected for subsequent PET experiments.

**Figure 1.**
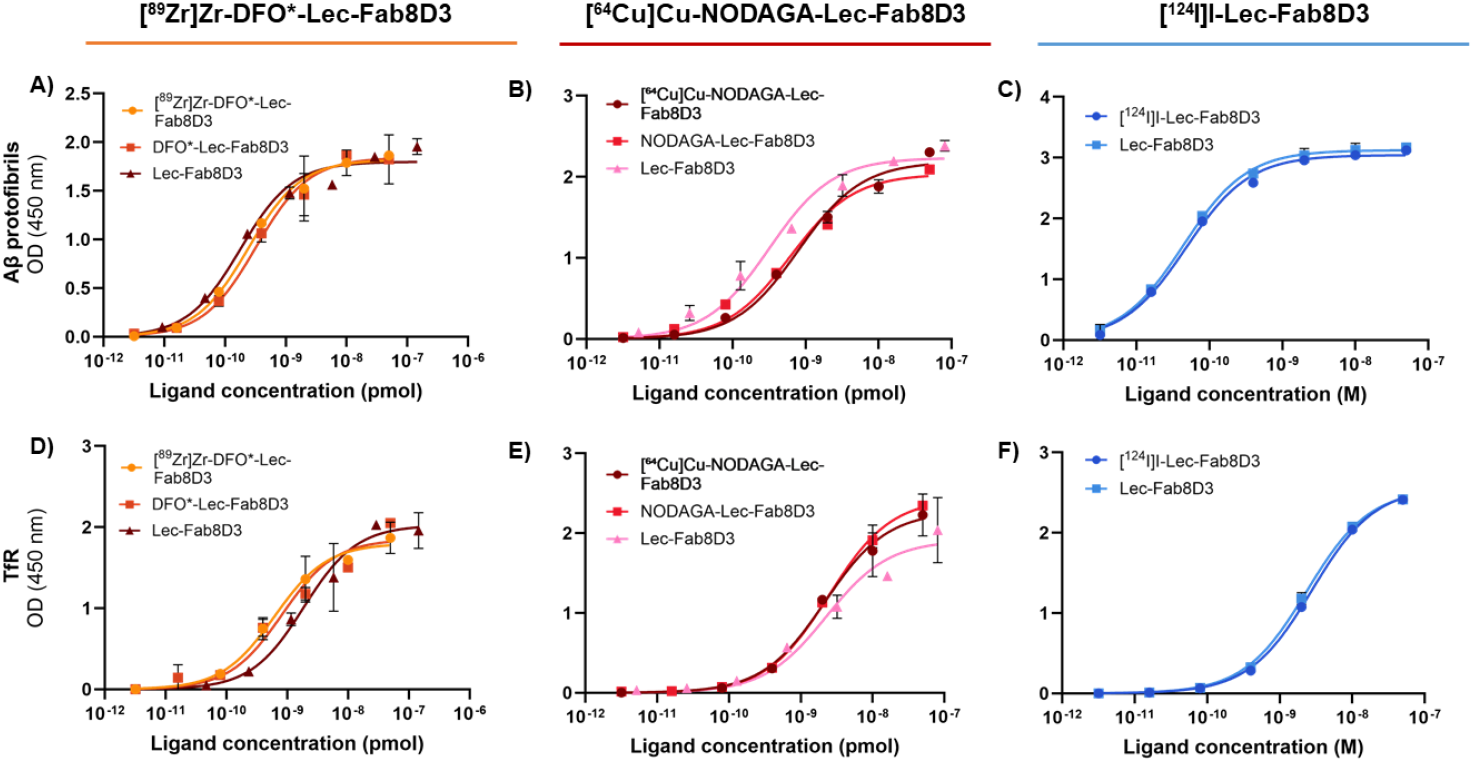
ELISA results for binding towards Aβ protofibrils of **A**) DFO*-Lec-Fab8D3 and radiolabeled [^89^Zr]Zr-DFO*-Lec-Fab8D3, **B**) NODAGA-Lec-Fab8D3 and radiolabeled [^64^Cu]Cu-NODAGA-Lec-Fab8D3 and **C**) [^124^I]I-Lec-Fab8D3 and towards mTfR of **D**) DFO*-Lec8D3-Fab8D3 and radiolabeled [^89^Zr]Zr-DFO*-Lec-Fab8D3, **E**) NODAGA-Lec-Fab8D3 and radiolabeled [^64^Cu]Cu-NODAGA-Lec-Fab8D3 and **F**) [124I]I-Lec-Fab8D3, showing no or minor alterations in antibody binding after modification and labeling.

Radiolabeling of Lec-Fab8D3 with [^89^Zr]Zr-oxalate, [^64^Cu]CuCl_2_, and [^124^I]NaI resulted in radiolabeled antibodies with high radiochemical purity (RCP) and radiochemical yield (RCY) for all three radiotracers (Table 2). The highest molar activity (Am) was achieved for [^64^Cu]Cu-NODAGA-Lec-Fab8D3 with 327 GBq/µmol at the end of synthesis. This high Am was particularly important in the case of ^64^Cu due to its shorter half-life compared to ^89^Zr and ^124^I and the therefore limited amount of radioactivity remaining at the PET scans acquired at 2-3 days post radiotracer administration. The binding capacity of the radiolabeled antibodies was assessed by ELISA and confirmed to be preserved toward both Aβ protofibrils and mTfR for all radiotracers (Figure 1).

**Table 2:** Summary of Lec-Fab8D3 radiolabeling results with ^89^Zr, ^64^Cu and ^124^I.

| Radiotracer | RCP (%) | RCY (%) | A<sub>m</sub> (GBq/$\mu\text{mol}$) | Final activity (MBq) |
| --- | --- | --- | --- | --- |
| [$^{89}\text{Zr}$]Zr-DFO*-Lec-Fab8D3 | 96 | 85 | 33.0 | 90 |
| [$^{64}\text{Cu}$]Cu-NODAGA-Lec-Fab8D3 | 96 | 95 | 327 | 723 |
| [$^{124}\text{I}$]I-Lec-Fab8D3 | 99 | 90 | 22.4 | 54 |

RCP = Radiochemical purity, RCY = Radiochemical yield, A<sub>m</sub> = Molar activity and final activity represents the radioactivity of collected labeled antibody after purification.

### PET imaging with ^89^Zr-, ^64^Cu-, and ^124^I-labeled Lec-Fab8D3 reveals Aβ pathology in Tg-ArcSwe mice

In vivo PET imaging was performed in 18-20-month-old Tg-ArcSwe mice and age-matched WT controls using Lec-Fab8D3 labeled with either [^89^Zr]Zr-oxalate (n=9), [^64^Cu]CuCl2 (n=9), or [^124^I]I (n=9). PET scans were acquired at 9 h, 24 h, and 72 h post-injection for the ^89^Zr and ^124^I groups. For the ^64^Cu group, imaging was conducted at 9 h, 24 h, and 48 h post-injection due to the shorter half-life of ^64^Cu (Figure 2, Figure S1). PET images representing radiotracer distribution in the brain at the last time point (48 or 72 h), clearly visualized Aβ-rich region in Tg-ArcSwe mice injected with radiolabeled Lec-Fab8D3 regardless of the isotope used, with the most intense signal seen in the cortex. The highest radioactivity was detected in Tg-ArcSwe mice that had received [^89^Zr]Zr-DFO*-Lec-Fab8D3 or [^64^Cu]Cu-NODAGA-Lec-Fab8D3. In contrast, the lowest radioactivity was found in WT mice administered with [^124^I]I-Lec-Fab8D3. Ex vivo autoradiography, providing a visual representation of radiotracer distribution in the absence of blood, further supported the in vivo PET results. Here, radiotracer accumulation was observed in the cortex, hippocampus and thalamus of Tg-ArcSwe mice. Interestingly, WT mice injected with radiometal (^64^Cu or ^89^Zr) labeled Lec-Fab8D3 displayed a faint but clear radioactive signal in the frontal cortex and central regions of the brain, despite absence of Aβ aggregates in WT mice. (Figure 2A). Autoradiography images also showed that Tg-ArcSwe mice had a varying degree of radioactivity accumulation in the cerebellum, consistent with lower cerebellar Aβ deposition in this mouse model (Figure 2B), on average 5-fold lower compared with the rest of the brain (Figure S2).^26^ To confirm antibody engagement with Aβ pathology in the brain, the administered Lec-Fab-8D3 was detected by anti-human IgG immunostaining which showed a close co-localization with Aβ pathology (Figure 2C).

**Figure 2.**
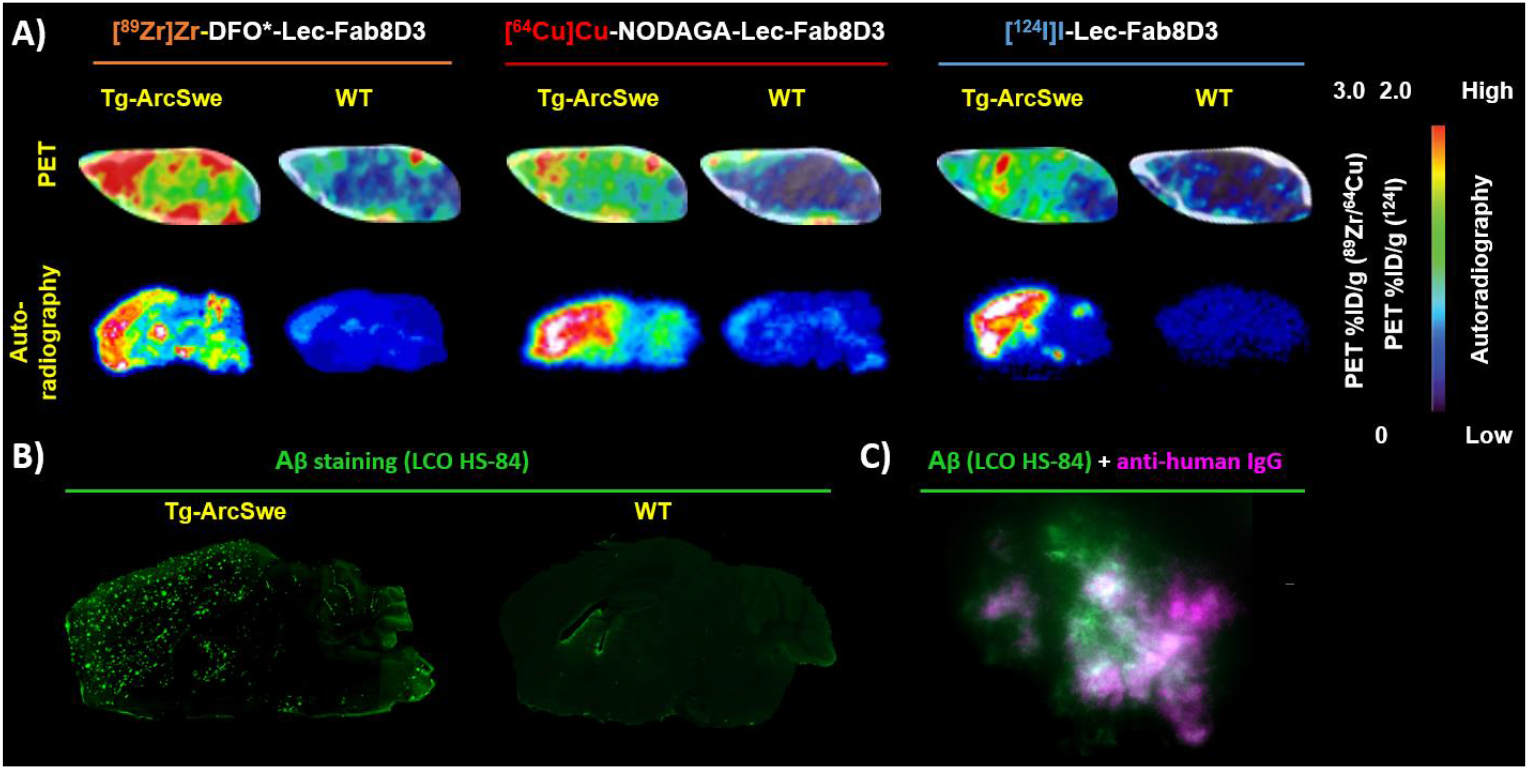
**A)** Representative sagittal PET images of the brain 2 or 3 days post-injection, showing cortical uptake of [^89^Zr]Zr-DFO*-Lec-Fab8D3 (left), [^64^Cu]Cu-NODAGA-Lec-Fab8D3 (middle), and [^124^I]I-Lec-Fab8D3 (right) in Tg-ArcSwe mice. Below PET images are corresponding autoradiography images of sagittal brain sections, confirming radiotracer accumulation in cortical regions associated with Aβ plaque deposition. Brain uptake is represented as percent of the injected dose per gram brain (%ID/g). **B)** Tg-ArcSwe and WT brain sections stained for Aβ pathology with the aggregate specific LCO stain HS-84. **C**) Combined Aβ (green; HS-84) and anti-human IgG staining (magenta) showing colocalization of the injected Lec-Fab8D3 tracer and Aβ plaques.

PET imaging at earlier time-points, i.e., 9 h and 24 h, provided a less distinct difference between the Tg-ArcSwe and WT mice due to the high concentration of radiolabeled antibody remaining in the bloodstream, which resulted in elevated radioactive signal across the entire brain region. (Figure S1).

### Antibody blood pharmacokinetics are unaffected by radiolabeling method

To compare the pharmacokinetic properties of the three antibody tracers, blood samples were collected from the time of administration of the radiolabeled antibody until the terminal end-point of the experiment. Blood concentrations (Figure 3A-C) measured over time revealed a similar clearance from the blood for all radiolabeled Lec-Fab8D3 variants with an average terminal blood concentration of 5.63 ± 0.01%ID/g, 7.18 ± 0.02%ID/g, 4.67 ± 0.02%ID/g for ^64^Cu, ^89^Zr and ^124^I radiolabeled antibodies, respectively, where it should be noted that the mice that had received [^64^Cu]Cu-NODAGA-Lec-Fab8D3 were sacrificed one day earlier than the mice that had received the two other variants of Lec-Fab8D3.

**Figure 3.**
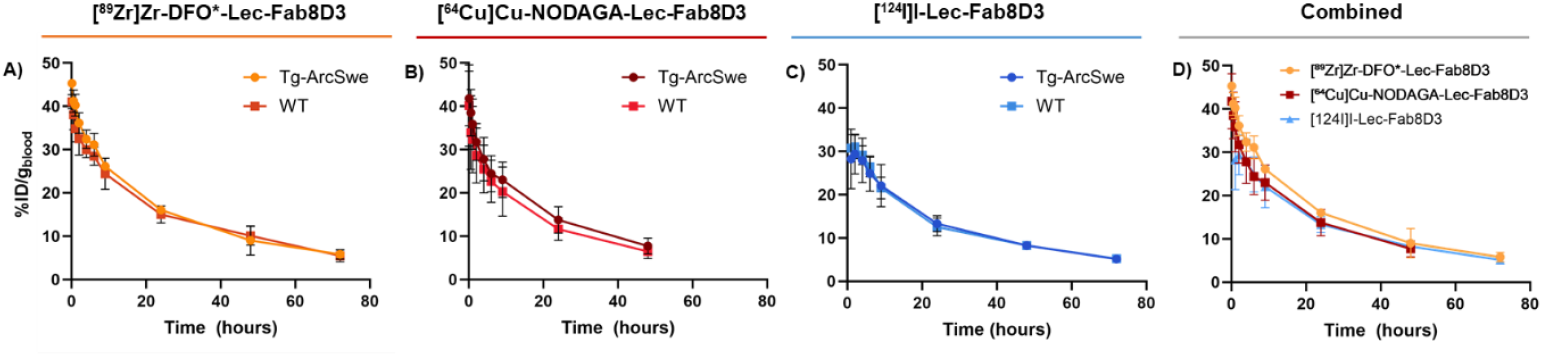
Blood elimination curves of **A**) [^89^Zr]Zr-DFO*-Lec-Fab8D3 (0-72 h); **B**) [^64^Cu]Cu-NODAGA-Lec-Fab8D3 (0-48 h); **C**) [124I]I-Lec-Fab8D3 (0-72 h) in Tg-ArcSwe and WT mice and D) all three radiotracers in Tg-ArcSwe mice

### PET quantification shows high Aβ-related brain retention and increasing brain-to-blood ratios over time

Quantification of radiotracer brain concentrations based on PET images showed no genotype difference in brain retention or brain-to-blood ratio at 9 h post-radiotracer administration (Figure 4A–F). With time, all radiotracers displayed an increased difference between Tg-ArcSwe and WT mice, both in terms of absolute brain concentration and when expressed as a brain-to-blood ratio. Specifically, [^89^Zr]Zr-DFO*-Lec-Fab8D3 and [^124^I]I-Lec-Fab8D3 were found at significantly higher concentration in the Tg-ArcSwe brain compared to the WT brain already at 24 h post-administration. As unbound tracer washed out of the brain, the difference between Tg-ArcSwe and WT mice increased to about 2-fold at 72 h. A similar trend was noted for [^64^Cu]Cu-NODAGA-Lec-Fab8D3, although statistical significance was not reached even at the last PET time point (48 h) due to a high variability between animals (Figure 4G–R). However, at the last PET imaging time point, all three radionuclides displayed significant differences between Tg-ArcSwe and WT mice in the brain-to-blood ratio, an important parameter for imaging contrast and specific PET signal. Notably, the radiometal-labeled antibodies exhibited higher overall brain concentrations in both Tg-ArcSwe and WT animals, compared to the radioiodinated antibody.

**Figure 4.**
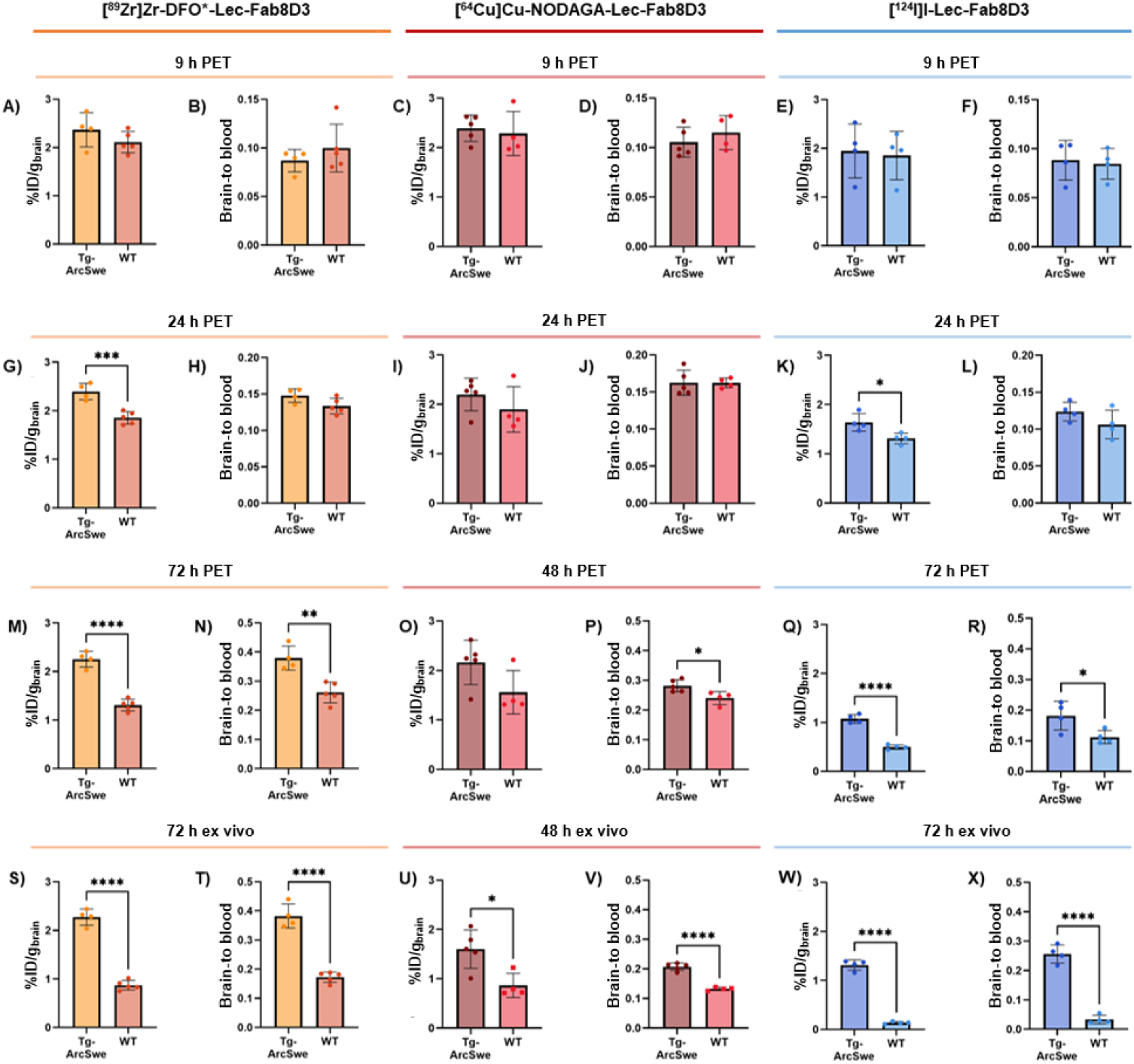
Quantification of brain concentration, expressed as % of injected dose per gram brain (%ID/g brain) and brain-to-blood ratio derived from PET imaging after 9 h (**A-F**), 24 h (**G-L**) and 48 h/72 h (**M-R**) after radiotracer administration and ex vivo gamma counting in perfused post-mortem brain tissue (**S-X**). Left (yellow/orange) represents [^89^Zr]Zr-DFO*-Lec-Fab8D3, Middle (purple/red) represents [^64^Cu]Cu-NODAGA-Lec-Fab8D3, and right (blue) represents [^124^I]I-Lec-Fab8D3.

### Ex vivo analysis confirms high Aβ-related brain retention and brain-to-blood ratios for all radionuclides

Ex vivo analysis, i.e. γ-counting, of perfused brains collected after the final PET scans further confirmed the increased brain uptake and brain-to-blood ratios in Tg-ArcSwe compared to WT mice for all three radionuclides at the terminal time points (Figure 4S–X). The most distinct differences in absolute brain concentrations were observed for [^89^Zr]Zr-DFO*-Lec-Fab8D3 and [^124^I]I-Lec-Fab8D3, while [^64^Cu]Cu-NODAGA-Lec-Fab8D3 showed a less pronounced but still detectable difference between Tg-ArcSwe and WT mice. Consistent with the in vivo PET results from the last scanning time point (Figure 3P), the variation between [^64^Cu]Cu-NODAGA-Lec-Fab8D3-administered animals was reduced after correction for blood concentration, resulting in a significant difference between Tg-ArcSwe and WT in brain-to-blood ratio (Figure 4V).

### Brain-to-cerebellum ratio discriminates between Tg-ArcSwe and WT only for radioiodinated antibody

PET quantification imaging is often simplified by calculating the ratio of radiotracer concentration between a region of interest and a reference region. As Tg-ArcSwe mice displayed lower Aβ levels in cerebellum, PET-derived brain-to-cerebellum ratios were calculated from the terminal PET-scans obtained for each antibody tracer (Figure 5). PET-based brain-to-cerebellum concentration ratios were approximately 1 in both genotypes for the radiometal-labeled [^89^Zr]Zr-DFO*-Lec-Fab8D3 and [^64^Cu]Cu-NODAGA-Lec-Fab8D3. In contrast, [^124^I]I-Lec-Fab8D3 showed a significantly higher ratio in Tg-ArcSwe compared to WT mice. Similarly, ex vivo brain-to-cerebellum ratios based on γ-counting were slightly above 1 for [^89^Zr]Zr-DFO*-Lec-Fab8D3 and [^64^Cu]Cu-NODAGA-Lec-Fab8D3, but with no genotype-dependent differences. Mice injected with [^124^I]I-Lec-Fab8D3 also exhibited elevated ratios, but again with a genotype difference, consistent with the PET derived brain-to-cerebellum ratio (Fig. 5C).

**Figure 5.**
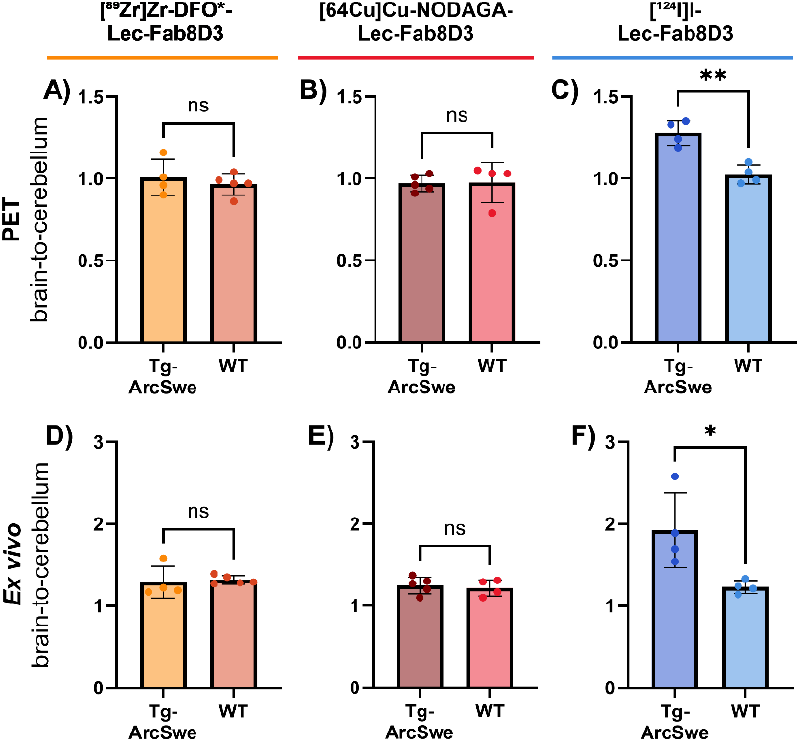
PET derived brain-to-cerebellum ratios in Tg-ArcSwe and WT mice **A**) 72 h after administration of [^89^Zr]Zr-DFO*-Lec-Fab8D3; **B**) 48 h after administration of [^64^Cu]Cu-NODAGA-Lec-Fab8D3; and **C**) 72 h after administration of [^124^I]I-Lec-Fab8D3. Corresponding ex vivo derived brain-to-cerebellum ratios for **D**) [^89^Zr]Zr-DFO*-Lec-Fab8D3; **E**) [^64^Cu]Cu-NODAGA-Lec-Fab8D3 and **F**) [^124^I]I-Lec-Fab8D3.

**Figure 6.**
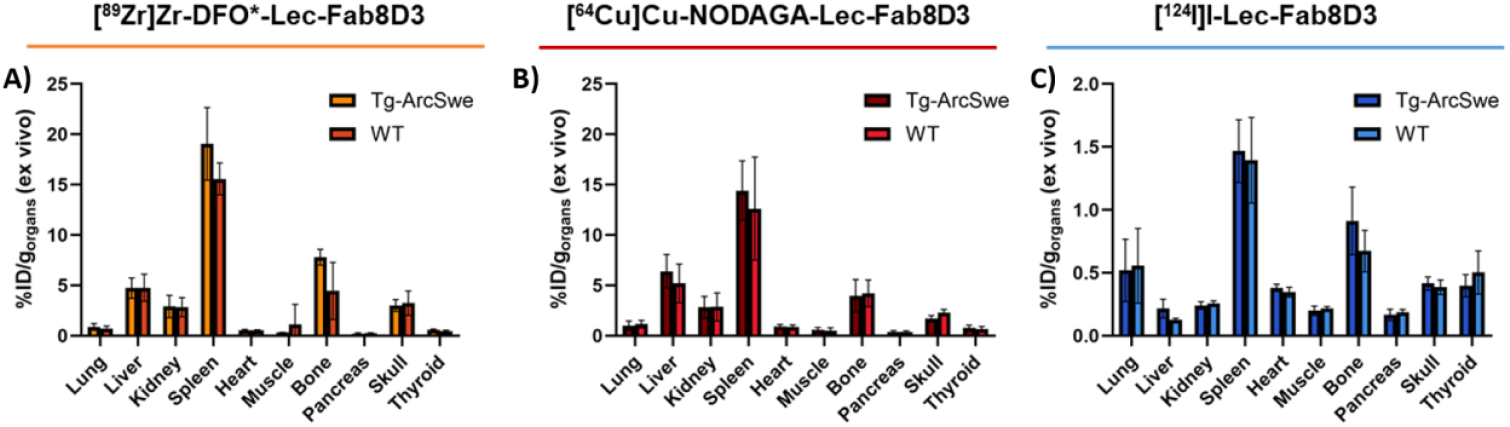
Ex vivo biodistribution to peripheral organs of **A**) [^89^Zr]Zr-DFO*-Lec-Fab8D3 (72 h p.i.), **B**) [^64^Cu]Cu-NODAGA-Lec-Fab8D3 (48 h p.i.) and **C**) [^124^I]I-Lec-Fab8D3 (72 h p.i.).

### Higher retention of radiometal-labeled Lec-Fab8D3 variants in peripheral organs

Terminal biodistribution to peripheral organs revealed higher overall concentrations for the radiometal-labeled antibodies compared to the radioiodinated variant. For all three radiolabeled variants of Lec-Fab8D3, the highest uptake was observed in the spleen, consistent with previous findings.^9,17,27,28^ TfR is highly expressed on reticulocytes, which are immature red blood cells released into circulation to mature. In rodents, the spleen plays a key role in reticulocyte release, contributing to the high uptake of TfR-targeting antibodies in this organ.^29^ Elevated signals were also detected in the bone and skull, likely due to antibody interaction with TfR expressed in the bone marrow. The relatively high liver signal observed with radiometal-labeled antibodies reflects progressive antibody degradation and intracellular accumulation of the radiometal.^30^ Similarly, degradation and deiodination of [^124^I]I-Lec-Fab8D3 resulted in a pronounced signal in the thyroid, where free iodine tends to accumulate.^24^

## Discussion

The development of immunoPET imaging of Aβ pathology presents a promising alternative to conventional amyloid PET imaging, as it may offer a more accurate representation of the Aβ species targeted by therapeutic antibodies. Given the complexity of antibody distribution and target engagement within the brain, the choice of radiolabeling strategy can significantly influence both the outcome and interpretation of Aβ immunoPET imaging. This study, therefore, aimed to compare PET imaging using Lec-8D3 labeled with three different radionuclides ^89^Zr, ^64^Cu, and ^124^I in a mouse model of Aβ pathology.

We first demonstrated successful chelator conjugation with DFO* and NODAGA, resulting in minor effects on affinity for Aβ and TfR. Subsequent radiolabeling using with [^89^Zr]Zr-oxalate, [^64^Cu]CuCl2 and [^124^I]I-NaI provided high radiochemical purity (RCP), radiochemical yield (RCY) and good molar activity (Am). For [^64^Cu]CuCl2 labeling, the Am was aimed higher than for the other radionuclides to compensate for the shorter half-life. TfR-mediated brain transport is dose-dependent and therefore administered similar antibody doses for all radiolabeled Lec-Fab8D3 variants, while maintaining sufficient injected radioactivity for multi-day PET imaging.^9,28^

PET imaging with Lec-Fab8D3 with the three radionuclides showed that the latest timepoints (48-72 h post radiotracer administration) gave the highest distinction in brain uptake and brain-to-blood ratio between Tg-ArcSwe and WT mice. This is due to the slow blood clearance of antibodies, with half-lives up to 30 days depending on its isotype and properties, but typically shorter for bispecific antibodies that carry a TfR binding domain, which can induce target mediated clearance.^17,31,32^

PET images and quantification showed higher overall brain concentrations of Lec-Fab8D3 radiolabeled with the metals ^89^Zr and ^64^Cu, compared to its radioiodinated version in both Tg-ArcSwe and WT mice, while all antibody variants displayed a similar absolute difference between the mouse genotypes. This phenomenon has been observed in previous studies of brain-targeted antibodies labeled with radiometals.^14,33–35^ It can likely be explained by TfR-mediated internalization of the bispecific antibody into cells, e.g. neurons, which are in high demand of iron and therefore express TfR on their surface. After internalization, the antibodies will be degraded but due to the residualizing properties of radiometals, the radiolabel will accumulate in the cell rather than being secreted, which will lead to a gradually increased non-target specific retention. Iodine, on the other hand, does not residualize and will therefore be excreted rapidly, reducing the overall retention^30,36^ Ex vivo gamma-counting data and autoradiography images strengthen this hypothesis further by revealing a higher retention in WT animals that received radiometal-labeled antibody compared to the WT mice administered with radioiodinated Lec-Fab8D3. This difference is particularly striking in the cortical region, where there is a high neuronal density and metabolic activity^37,38,37^ The residualizing effect of radiometals compared to iodine was also obvious in the ex vivo biodistribution to peripheral organs, where substantially higher overall radioactive signal was recorded in animals that received radiometal-labeled Lec-Fab8D3 than in animals that were given the radioiodinated Lec-Fab8D3. Interestingly, while [^89^Zr]Zr-DFO*-Lec-Fab8D3 remained stable in Tg-ArcSwe mice over time, [^64^Cu]Cu-NODAGA-Lec-Fab8D3 showed a subtle decline between 24 h and 48 h. This could suggest that even though NOTA/NODAGA is a suitable choice of chelator for 64Cu-labeling it may be less stable than the 89Zr-DFO* complex in vivo, which could make ^64^Cu more prone to be secreted after intracellular degradation.

Radiometal accumulation in brain regions lacking the antibody target can lead to increased background signal, reduced contrast, and impaired detection of low-abundance targets. This hampers the ability to distinguish between brain regions with varying target expression. This effect is evident when comparing brain-to-cerebellum ratios in mice imaged with different radionuclides in this study. Consistent with previous studies on aged Tg-ArcSwe mice, post-mortem analysis confirmed approximately fivefold higher Aβ concentrations in the brain compared to the cerebellum. Accordingly, higher antibody binding and retention are expected in the brain. This was observed with [^124^I]I-Lec-Fab8D3, which showed significantly higher PET-derived brain-to-cerebellum ratios in Tg-ArcSwe than in WT mice. In contrast, both radiometal-labeled antibodies yielded ratios near unity, with no genotype-dependent differences. One possible explanation is that the blood component of the brain may obscure regional differences, particularly since the cerebellum may have relatively higher perfusion. Supporting this, ex vivo brain-to-cerebellum ratios and autoradiography, which both exclude the blood signal, revealed higher cortical uptake in Tg-ArcSwe mice injected with [^89^Zr]Zr-Lec-Fab8D3, reflecting Aβ accumulation. However, there was still no difference between the genotypes, as a similarly increased signal was seen in the cortex of WT animals, in this case likely due to antibody accumulation in neurons.

Thus, although the radiotracers carrying ^64^Cu or ^89^Zr showed the ability to clearly distinguish between Tg-ArcSwe and WT and yielded a higher overall signal compared with radioiodine, the discussion above illustrates the difficulty to balance the dual targets of the bispecific antibodies. Given that Lec-Fab8D3 has 10-100-fold higher affinity for Aβ over TfR, the signal derived from Aβ rich brain areas likely reflects true target binding. In contrast, regions high in TfR expression but lacking Aβ may accumulate a considerable amount of radiometal signal, especially given the long accumulated time of potential TfR-mediated intracellular antibody uptake. In practice, most brain regions express both Aβ and TfR to varying degrees, complicating efforts to disentangle their respective contributions to the observed signal.

All three investigated radionuclides – ^89^Zr, ^64^Cu and ^24^I – have been used in preclinical immunoPET setting for targets within the brain, but have not been translated into a clinical setting or compared systematically for brain PET.^13,14,38–41^ This work shows that Lec-Fab8D3 labeled with either ^89^Zr, ^64^Cu or ^124^I allows imaging of Aβ in the AD mouse brain but that the choice of isotope has to be carefully considered. The slow pharmacokinetics of antibodies makes ^64^Cu less suitable than ^89^Zr and ^124^I. Both ^89^Zr and ^124^I appeared to be highly suitable for immunoPET, making the choice of radionuclide mainly dependent on other factors e.g. availability and dosimetry. In addition, while ^124^I gives a more clean picture of Aβ pathology, ^89^Zr provides additional potentially important information about the accumulated off-target antibody distribution in the brain.

In future work, imaging with ^89^Zr and ^124^I could be considered at later timepoints, e.g. 7 days, to have less radiolabeled antibody remaining in the blood. It would also be of interest to evaluate radiolabeled Lec-Fab8D3 in younger mice to examine this radiotracer as a potential early diagnostic marker for AD and to develop bispecific Lec in different antibody formats and sizes to reduce the biological half-life. Moreover, engineering Lec-Fab8D3 with a human TfR (hTfR) binder instead of a murine TfR (mTfR) binder would allow for clinical evaluation and potential clinical translation.

## Conclusions

ImmunoPET imaging with Lec-Fab8D3 targeting Aβ in AD offers significant opportunities for earlier and more specific detection of pathology and may serve as a companion diagnostic to current and emerging immunotherapies. Among the radionuclides tested, ^89^Zr and ^124^I proved to be the most promising isotopes for PET imaging with radiolabeled Lec-Fab8D3, due to their suitable half-lives and strong PET signal following amyloid pathology in the AD mouse brain. Future work will focus on imaging at later timepoints with ^89^Zr and ^124^I, decreasing the biological half-life of our antibody, and further optimizing PET contrast to support early diagnostic applications.

## Supporting information

Lopes vd Broek, Supplementary information

## Abbreviations

%ID/g: percent of injected dose per gram
Aβ: amyloid beta
AD: Alzheimer’s disease
Am: molar activity
BBB: blood-brain barrier
CT: computed tomography
DFO: deferoxamine
Fab: fragment antigen-binding fragment
IHC: immunohistochemistry
iTLC: instant thin layer chromatography
Lec: Lec
mTfR: murine transferrin receptor
NCS: N-chlorosuccinimide
PET: positron emission tomography
RCP: radiochemical purity
RCY: radiochemical yield

## Materials and Methods

### Expression and purification of Lec-Fab8D3

Lec-Fab8D3 was based on the humanized monoclonal antibody Lecanemab that binds selectively to aggregated forms of Aβ.^42,43^ The bispecific, brain penetrating Lec-Fab8D3 was designed based on a previously published format^17,44^, with a single Fab fragment of the TfR-binding antibody 8D3^45^ attached to the C-terminus of one of the antibody’s heavy chains, using the knob-into-hole technique. The mutations (L234A, L235A, P329G) were introduced to the Fc-domain to reduce effector functions.46 The antibody was transiently expressed in Expi293F cells and purified using protein A chromatography, followed by size-exclusion chromatography, according to a previously described protocol.^47^

### ELISA

**Coating**. The 96-well half-area plates (Corning Inc., New York, NY) were coated overnight at 4°C with Aβ protofibrils (50nM diluted in PBS) mTfR (50 µL/well, 5 µg/mL diluted in PBS), or hIgG (0.5 µL/well, 5 µg/mL diluted in PBS).**Blocking**. The wells were blocked with 1% BSA in PBS for 1h. The proteins were diluted in incubation buffer (PBS containing 0.1% BSA, 0.05% Tween-20 and 0.15% Kathon) **Primary antibody**. The plates were washed with wash buffer (phosphate buffer with 0.1% Tween-20 and 0.15% Kathon, pH 7.5). Lec-Fab8D3 their radiolabeled equivalents were serial diluted in incubation buffer (PBS with 0.1% BSA, 0.05% Tween-20 and 0.15% Kathon) from 50 nM to 3.2 pM and incubated for 2 hours. **Secondary antibody**. The plates were washed with wash buffer and detected with horseradish peroxidase (HRP)-coupled goat anti-human IgG-F(ab’)2 (1:2000, Jackson ImmunoResearch Laboratories, West Grove, PA, 109-036-006). **Plate development**. Signals were developed with K blue aqueous TMB substrate (Neogen Corp., Lexington, KY), quenched with 1M H2SO4 and read with a spectrophotometer at 450 nm. The data was analyzed using Excel and GraphPad Prism.

### Animals

The Tg-ArcSwe mouse model harbours the Arctic (AβPP E693G) and Swedish (AβPP KM670/671NL) mutations, is maintained on a C57BL/6 background and shows elevated levels of soluble Aβ protofibrils at a young age and abundant and rapidly developing plaque pathology starting at around 6 months of age. The fibrils of the Aβ aggregates show high similar to those seen in human sporadic AD [REF Zielinski.^48^ Both Tg-ArcSe males (n=X) and females (n=y) were used and littermates (males, n=y; females, n=z) were used as control animals (WT). The animals were housed in rooms with controlled temperature and humidity in an approved facility at Uppsala University with *ad libitum* access to food and water. All procedures described in this paper were approved by the Uppsala County Animal Ethics board (5.8.18-16493/2024), following the rules and regulations of the Swedish Animal Welfare Agency and in compliance with the European Communities Council Directive of 22 September 2010 (2010/63/EU).

### DFO*/NODAGA conjugation

Lec-Fab8D3LALA was conjugated with either DFO*-NCS (ABX, 7272) or p-NCS-benzyl-NODA-GA (Chematech, C103) for radiolabeling with ^89^Zr and ^64^Cu, respectively. Prior to conjugation, the antibody was buffer-exchanged into 0.1 M metal-free NaHCO_3_/Na_2_CO_3_ buffer (pH 9.0) using 7 kDa MWCO ZebaSpin desalting columns (Thermo Fisher). The final antibody concentration was adjusted to 4.5 mg/mL, with a total antibody amount of 500–1000 μg. The chelators were prepared as 5 mg/mL for DFO*-NCS or 2,5 mg/mL for p-NCS-benzyl-NODA-GA solutions in DMSO and added to the antibody solution at a molar excess of 7-fold for DFO-NCS* and 10-fold for NODAGA-NCS. The conjugation reaction was carried out for 2 hours at 37°C under gentle mixing. Following incubation, excess unreacted chelator was removed by purification with 7 kDa MWCO ZebaSpin columns, and the conjugated antibody was collected in metal-free 0.9% NaCl and stored at 4°C until radiolabeling. The final antibody concentration was determined using a Spectrophotometer (DeNovix DS-11).

### Radiolabeling with ^89^Zr

[^89^Zr]Zr-oxalate in 1M oxalic acid was provided by Karolinska Institutet. 40 µL of oxalic acid (1M) was added to 60 µL of [^89^Zr]Zr-oxalate (106 MBq) and 45 µL of Na_2_CO_3_ (2M) was added slowly in a stepwise manner. The pH was adjusted using oxalic acid (1M) until pH 4.5 was reached and the mixture was incubated for 3 min at room temperature. 214 µL of metal-free HEPES buffer (0.5M, pH 7) was added, followed by 120 µL of DFO*-Lec-Fab8D3LALA (550 µg, 4.57 mg/mL). The reaction mixture was incubated at 37°C for 2 hours. Radiolabeled antibody was purified using a NAP-5 column (Cytiva) and eluted in three fractions (150 µL, 700 µL, and 150 µL) in PBS, yielding 90 MBq of purified [^89^Zr]Zr-DFO*-Lec-Fab8D3LALA, with a RCP>96%, RCY=85% and a a Am of 33.0 GBq/µmol. RCP was assessed using radio-TLC on iTLC-SG paper (Agilent, SGI0001) with 20 mM citric acid + 60 mM EDTA : acetonitrile (9:1) buffer as the mobile phase. The retention factors (Rf) were: [^89^Zr]Zr-DFO*-Lec-Fab8D3: Rf = 0, unbound [^89^Zr] DFO*-NCS: Rf =0,6, Free [^89^Zr]Zr^4+^:Rf = 1

### Radiolabeling with ^64^Cu

[^64^Cu]CuCl_2_ was purchased from Risø, Roskilde, DTU, Denmark. Prior to radiolabeling, NODAGA-Lec-Fab8D3LALA was buffer-exchanged into metal-free NaOAc buffer (0.1M, pH 5.5), using a 7 kDa MWCO ZebaSpin desalting column (Thermo Fisher). For radiolabeling, [^64^Cu]CuCl_2_ was reconstituted in NaOAc (0.1M, pH 5.5), and 40 μL [^64^Cu]CuCl_2_ (824 MBq) was added to 115 μL NODAGA-antibody solution (444 μg, 3.85 mg/mL). The reaction was incubated at for 45 min at 37°C, after which additional NaOAc buffer (0.1M, pH 5.5) was added to a final reaction volume of 500 μL. The labeled antibody was purified using a NAP-5 size-exclusion column (Cytiva) and eluted in 1 mL PBS, yielding 723 MBq of [^64^Cu]Cu-NODAGA-Lec-Fab8D3LALA with a RCP>96%, RCY of 94% and a Am of 327 GBq/μmol (EOS). RCP was assessed using radio-TLC on iTLC-SG paper (Agilent, SGI0001) with 20 mM citric acid + 60 mM EDTA : acetonitrile (9:1) buffer as the mobile phase. The retention factors (Rf) were: [^64^Cu]Cu-NODAGA-Lec-Fab8D3: Rf = 0, unbound [^64^Cu]Cu-NODAGA-NCS: Rf = 0,5, Free [^64^Cu]CuCl_2_: Rf = 1

### Radiolabeling with ^124^I

For ^124^I-labelling, 130 µl [^124^I]NaI solution (Advanced Center Oncology Macerata, Montecosaro, Italy) was pre-incubated 15 min with 13 µl NaI (100 µM) before addition of 70,2 µL of Lec-Fab8D3 (420 µL, 5,98 mg/mL) and 4,8 µL Chloramine-T (4 mg/mL) in PBS, mixed in PBS to a final volume of 273 µl. After 120 s the reaction was quenched by addition of 9,7 µL of sodium metabisulfite (4 mg/mL) in PBS. Radiolabeled antibody was purified using a NAP-5 column (Cytiva) and eluted and eluted in 1 ml of PBS to yield 54 MBq of [^124^I]I-Lec-Fab8D3 with a RCP>99%, RCY of 90% and a Am of 22.4 GBq/μmol (EOS). RCP was assessed using radio-TLC on iTLC-SG paper (Agilent, SGI0001) with 70% acetone in water as the mobile phase. The retention factors (Rf) were: [^124^I]I-Lec-Fab8D3: Rf = 0, Free [^124^I]Na:I Rf = 1

### PET imaging

Animals belonging to the [^124^I]Lec-Fab8D3LALA group were given water supplemented with 0.2% NaI on the day before radiotracer administration to reduce thyroidal uptake of ^124^I. Mice were were anesthetized using 5% sevoflurane and i.v. injected with 4.92 ± 0.71 MBq [^89^Zr]Zr-DFO*-Lec-Fab8D3 (n=9), 31.23 ± 5.26 MBq [^64^Cu]Cu-NODAGA-Lec-Fab8D3 (n=15) or 4.68 ± 0.72 MBq [^124^I]I-Lec-Fab8D3 (n=9), corresponding to 0.86 ± 0.11, 0.55 ± 0.02 and 1.20 ± 0.07 µg antibody / gram mouse respectively. During the PET scan, animals were anesthetized using 2-3% sevoflurane in medical air supplemented with oxygen gas (0.2 L/min). Three animals were placed in the scanner gantry – two side by side, one on top - with the head and chest in the field of view and the mice were scanned 9 hours, 24 hours, 48 hours or 72 hours after radiotracer administration, with a maximum of three scans per animal. Dynamic PET images were acquired for 30 min for the 9 hour and 24 hour post injection PET scans, and 60 min for the 48 hour post injection and 72 hour post injection PET scans using a PET/MRI scanner (Nanoscan PET-3T MRI, Mediso, Medical Imaging Systems, Budapest, Hungary). Animals with [^64^Cu]Cu-NODAGA-Lec-Fab8D3 were scanned for 2 hours for the 72 hour post administration PET scan. After PET scans, the chamber bed was transferred to a SPECT/CT scanner (Nanoscan SPECT (4H)-CT, Mediso, Medical Imaging Systems, Budapest, Hungary), and the animals underwent a 6 min 80-kV CT scan.

PET data was reconstructed into 10 minute frames using the ordered subsets expectation-maximization (OSEM) 3D algorithm and CT raw files were reconstructed using Filter Back Projection (FBP). All subsequent processing of the PET and CT images was performed in imaging software Amide 1.0.6.^49^ The CT scan was manually aligned with an MRI based mouse brain atlas^50^ containing outlined regions of interests e.g. hippocampus, striatum, thalamus, cerebral cortex, prefrontal cortex and cerebellum. The PET image was then aligned with the CT and MRI-atlas and PET data was quantified. In all PET experiments, mice were scanned in a randomized order.

### Ex vivo biodistribution

After 2, 48 or 72h, mice were anaesthetized with isoflurane and a terminal blood sample was taken from the heart, followed by transcardial perfusion with 40 mL of 0.9% NaCl for 2.5 to clear the brain and organs from blood. Thereafter, lung, liver, kidney, whole heart, pancreas, spleen, femoral muscle, femoral bone, skull bone and submandibular glands were isolated to evaluate the biodistribution of the radiolabeled compounds. The brain was divided into left and right hemispheres, and the left hemisphere was further divided into cerebrum and cerebellum. The brain samples were immediately frozen on dry ice. The radioactivity of the samples was measured with a gamma-counter (2480 Wizard™, Wallac Oy PerkinElmer, Turku, Finland). Protein concentrations were expressed as percent of injected dose per gram tissue (%ID/g).

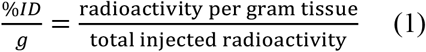

The brain-to-blood ratio was calculated by dividing the radioactivity in the brain per gram brain and the radioactivity in the blood per gram blood.

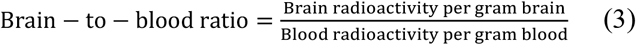

### Autoradiography

Upon PET scanning and transcardial perfusion, the spatial distribution of the three radiolabeled Lec-Fab8D3 variants was analyzed with autoradiography. Right hemispheres were cryosectioned sagittally with a thickness of 20 µm and sections were exposed together with a radioactive standard to phosphor imaging plates (PerkinElmer). After 3-7 days, the phosphor plate was scanned with an Amersham Typhoon imager (Cytiva) with a 50 µm resolution. Images were processed in ImageJ.

### Immunofluorescence

Immunofluorescence was performed to visualize Aβ pathology and localize the administered antibody in sagittal cryosections of mouse brain tissue. Tissue sections (20 µm thick) were initially fixed in ice-cold methanol for 10 minutes and subsequently rinsed twice in phosphate-buffered saline (PBS) for 5 minutes each. To block unspecific binding and permeabilize the tissue, sections were incubated for 1.5 hours in a blocking solution containing 5% normal goat serum (NGS) and 0.4% Triton X-100 in PBS. After blocking, sections were washed three times for 5 minutes in PBS. Antibody staining was performed by incubating the sections with goat anti-human Alexa Fluor 647-conjugated antibody (109-606-006, NordicBiosite) diluted 1:200 in PBS 0.05% Tween-20 for 2 hours at room temperature. Following this incubation, slides were washed twice for 5 minutes in PBS. To label Aβ fibrils, sections were then incubated with the luminescent conjugated oligothiophene (LCO) HS-84, diluted to a final concentration of 50 nM in PBS0.05% Tween-20, for 15 minutes at room temperature under gentle shaking. Excess dye was removed by washing the slides three times for 5 minutes in PBS. Sections were mounted using a mounting medium containing DAPI and imaged using a Zeiss Observer Z.1 microscope with ZEN 3.7 software (Carl Zeiss Microimaging GmbH, Jena, Germany).

### Aβ quantification in brain tissue

The left hemispheres from PET scanned animals were divided into brain (cerebrum) and cerebellum, and frozen at -80, then homogenized at a 1:5 tissue:buffer ratio in tris buffered saline supplemented with complete protein inhibitor cocktail using a Precellys Evolution homogenizer (Bertin Technologies) (4×10 s at 5500 rpm). Homogenates were centrifuged 1 h at 16 000xg and the supernatant was removed, while the pellet was resuspended and homogenized with 70% formic acid, to dissolve fibrillar Aβ. After 1 h centrifugation at 16 000xg, the supernatant was collected, neutralized with 2 M Tris and diluted (1:10 000) in MSD sample diluent, then analyzed for total Aβ38, Aβ40 and Aβ42 with the V-PLEX® Aβ peptide panel 1 (6E10) immunoassay (Meso Scale Discovery, K15200E), according to the manufacturer’s instructions.

### Statistical analyses

Statistical analyses were performed in GraphPad Prism 10.1.0 (GraphPad Software, Inc., San Diego, CA). The results are reported as mean ± standard deviation and statistical assessment was conducted by unpaired t-test or a two-way ANOVA with multiple comparisons test.

## Acknowledgements

We would like to thank Lara Garcia-Varela, University of Santiago de Compostela for her contributions to introduce ^89^Zr-labeling and Lars Nilsson, Oslo University, for the development of the Tg-ArcSwe model. The department of nuclear medicine and medical physics at Karolinska University hospital is also greatly acknowledged for providing the ^89^Zr-isotope. The molecular imaging work in this study was performed at the Preclinical PET-MRI Platform, a research infrastructure at Uppsala University, Sweden.

