## Supplementary material for "Radionuclide selection influences imaging outcomes in immunoPET with a brain-penetrant anti-Aβ antibody": Lopes vd Broek, Supplementary information

**SUPPLEMENTAL FIGURES**

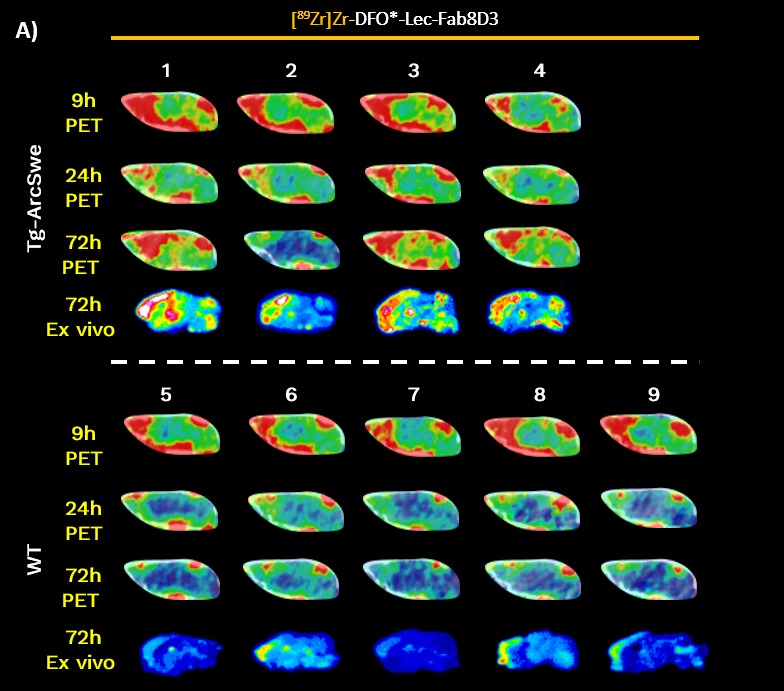

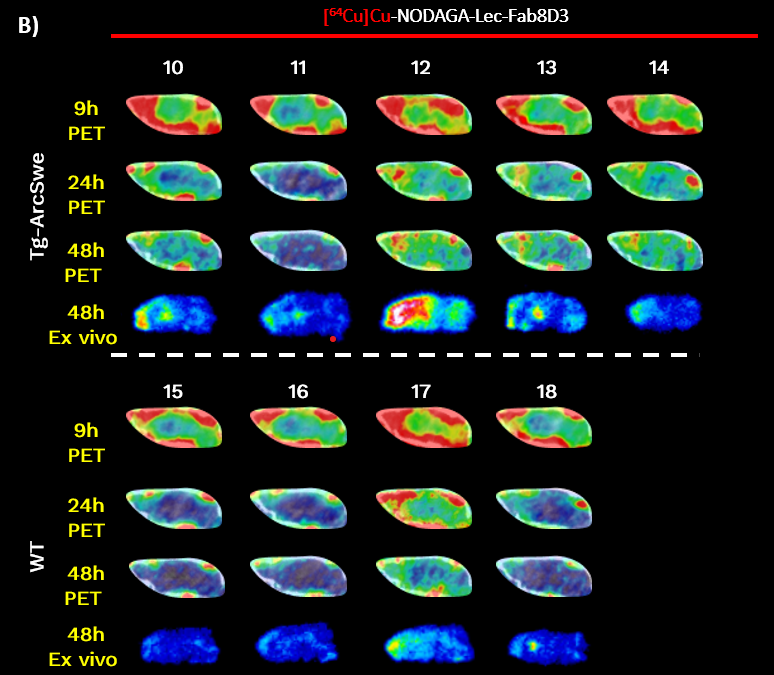

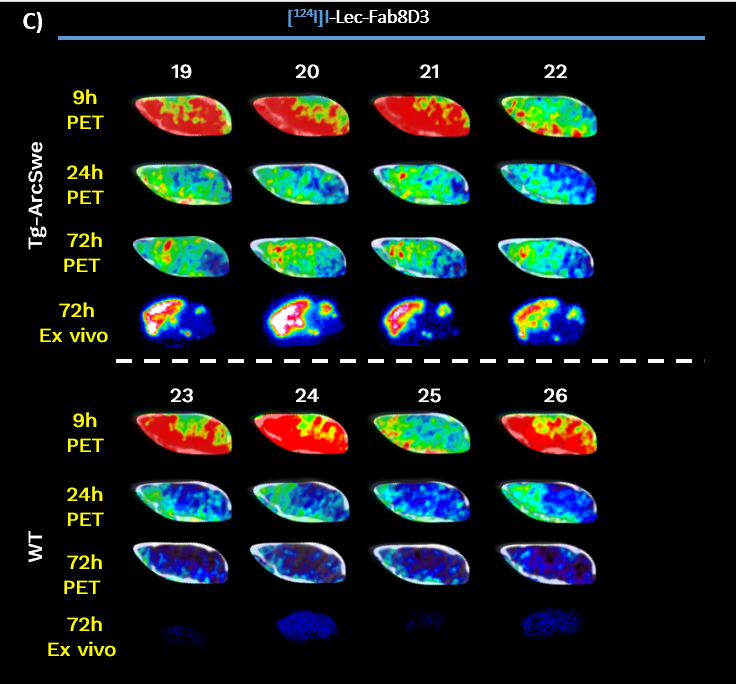

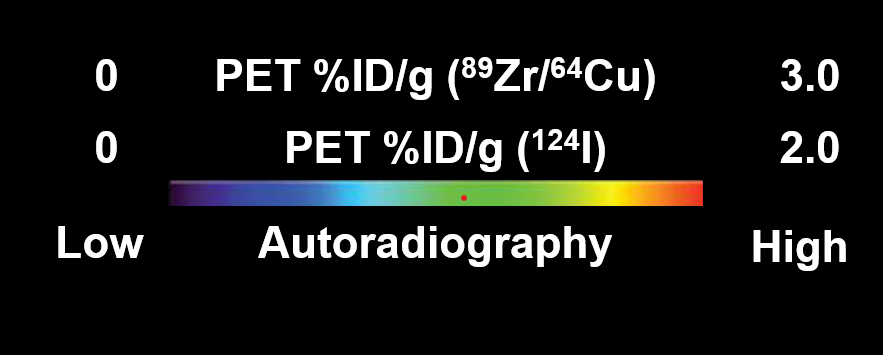

**Figure S1. PET and** **ex vivo brain images at 9, 24, 48 and 72 h post-injection.** Individual PET images of the brain 9, 24, 48 and 72 hours post-injection, showing cortical uptake of [⁸⁹Zr]Zr-DFO*-Lec-Fab8D3 (A), [⁶⁴Cu]Cu-NODAGA-Lec-Fab8D3 (B), and [¹²⁴I]I-Lec-Fab8D3 (C) in Tg-ArcSwe mice. Below PET images are corresponding autoradiography images of sagittal brain sections, confirming radiotracer accumulation in cortical regions associated with Aβ plaque deposition. Brain uptake is represented as percent of the injected dose per gram brain (%ID/g).

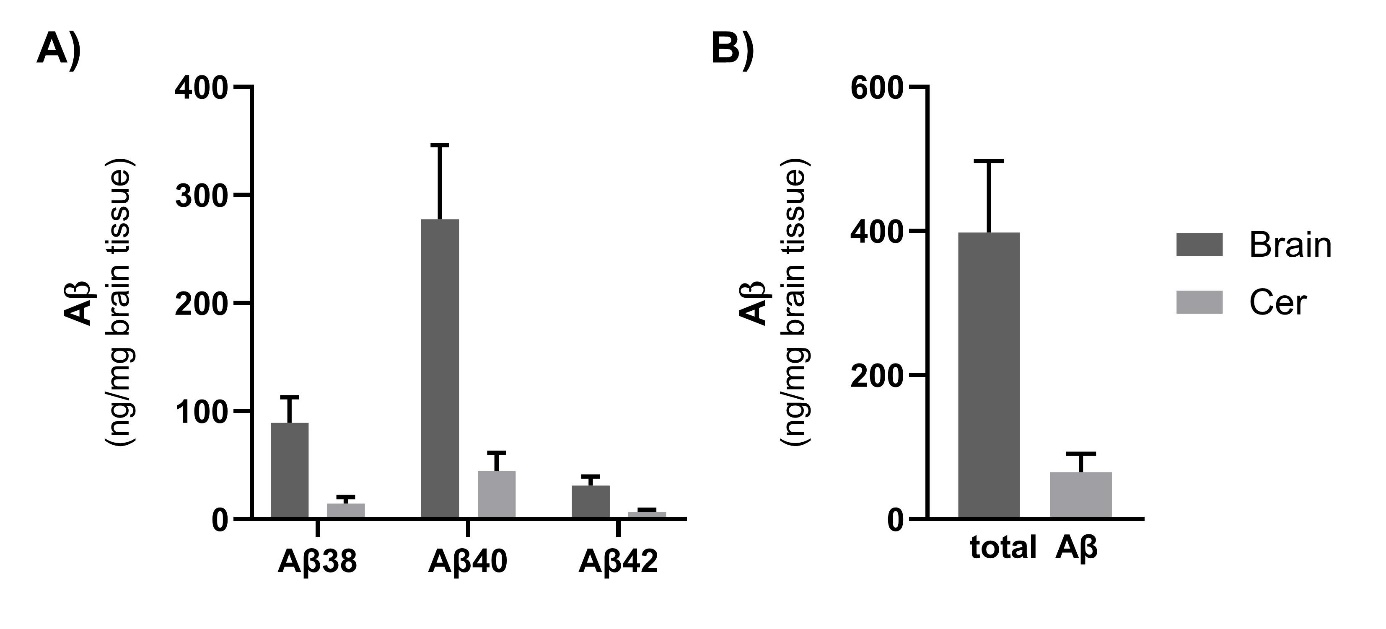

**Figure S2. Aβ quantification in mouse brain tissue. A)** Quantification of Aβ38, Aβ40 and Aβ42 in brain tissue extracts from all PET scanned mice determined by MSD immunoassay. **B**) Total Aβ (sum of Aβ38, Aβ40 and Aβ42 in **A**) in brain tissue extracts from all PET scanned mice.
